# Incorporating environmental heterogeneity and observation effort to predict host distribution and viral spillover from a bat reservoir

**DOI:** 10.1101/2023.04.04.535562

**Authors:** Rita Ribeiro, Jason Matthiopoulos, Finn Lindgren, Carlos Tello, Carlos M. Zariquiey, William Valderrama, Tonie E. Rocke, Daniel G. Streicker

## Abstract

Predicting the spatial occurrence of wildlife is a major challenge for ecology and management. In Latin America, knowledge of the true number and locations of vampire bat colonies precludes informed allocation of measures intended to limit lethal rabies spillover to humans and livestock. We inferred the complete spatial distribution of vampire bat roosts while accounting for observation effort and environmental covariates by fitting a Log Gaussian Cox Process model to the locations of 563 roosts in three regions of Peru. Our model explained 47% of the variance in the observed roost distribution and identified landscape correlates of roost establishment. Our model estimated that 1,795 roosts (76%) remain undiscovered and identified hotspots of undetected roosts in currently rabies-free areas, implying high risk for viral incursion. Incorporating the locations of undetected roosts improved spatial predictions of rabies spillover to livestock, revealed areas with disproportionate underreporting to surveillance systems, and indicated a higher rabies burden than previously estimated. We provide a robust approach to infer the distribution of a mostly unobserved bat reservoir that can inform strategies to prevent the re-emergence of an important zoonosis.

## Introduction

Mapping the geographic distribution of animal reservoirs of infection is a pre-requisite to anticipate and prevent spillover to other species [1]. For example, vaccination campaigns for dog rabies use detailed surveys of dog density to allocate vaccine distributions, and control programmes of bovine, sheep, and goat brucellosis require livestock census for adequate implementation of testing, vaccination, and slaughter strategies [2–4]. However, for wild or free-ranging species, distribution data are frequently sparse, presence-only records that do not cover the full geographic range [5,6]. The problem is exacerbated for species such as bats and rodents that are key reservoirs for zoonoses, but are difficult to observe given their small body size, reclusive nature, nocturnal activity, or high mobility [5,7,8]. Species distribution models (SDMs) that use relationships between field observations and environmental data to predict the occurrence or abundance of disease vectors or reservoirs are widely used to anticipate disease spread and help policymakers target prevention and control strategies [9]. More recent extensions of SDMs can deal with observation bias in input data and explicitly account for autocorrelation arising from geographic proximity between observations, hence capturing residual autocorrelation in the data, but have only rarely been applied to wildlife disease reservoirs. Accounting for such inferential challenges is vital to avoid misguided spatial allocations of interventions [5,6,10,11].

One of the most important zoonotic viruses transmitted by bats is rabies (*Rhabdoviridae lyssavirus*), which is capable to cause acute lethal encephalitis in all mammals [12]. In Latin America, the common vampire bat (*Desmodus rotundus*) is the main reservoir of rabies due to its population abundance, wide geographic distribution and obligate blood feeding, which provides a direct route for human and livestock infection via contaminated saliva [13,14]. Efforts to mitigate the burden of vampire bat rabies (VBR) include human and livestock vaccination; however, financial and logistical challenges result in campaigns that are largely reactive to rabies mortality events, and costs in human lives and livestock persist [15–17]. VBR management also involves controlling vampire bat populations using topical anticoagulant poisons that spread between individual bats during social grooming or that are applied to cattle for later consumption by bats [18,19]. The ability of bat culling to reduce rabies incidence is controversial, and efficacy has been questioned based on field, phylogenetic, and modelling studies [15,20–22]. Further, countries with active culling campaigns have seen viral spatial expansions and growing disease burdens [23,24]. It is hypothesized that culls may be ineffective because rabies virus is maintained by spatial processes, including wave-like invasions into historically rabies-free areas and metapopulation maintenance among patchworks of bat colonies, but culls are reactive to rabies outbreaks and rarely synchronized across enzootic regions [21,23]. Improving the implementation of culls or the spatial distribution of human and livestock vaccines requires knowledge of the spatial distribution of vampire bat populations and how this distribution determines spillover risk; however, with rare exceptions, neither the location nor the number of vampire bat roosts are known completely [25].

Previous models have predicted the distribution of *D. rotundus* over large geographic scales (e.g., across countries) and have explored how climate change might affect the future distribution of this species [26–28]. Predictions from these models are necessarily coarse given the wide variety of habitat types considered. In contrast, informing on-the-ground management requires high resolution predictions that are robust to spatial autocorrelation and that explicitly consider heterogeneities in the search effort applied to detect individuals. Here, we used data on the locations of *D. rotundus* roosts (any place a wild bat uses for shelter or protection) to predict the distribution and expected density of detected and undetected roosts while accounting for spatial correlation structures and heterogeneities in observation effort that generated the underlying roost data. Roost data were collected in three regions in southern Peru where vampire bat rabies virus (VBRV) is either endemic or invading historically uninfected zones. Our specific goals were to (1) identify the environmental and anthropogenic variables that drive roost distribution; (2) reconstruct the map of expected *D. rotundus* roost density, controlling for heterogeneous effort; (3) estimate the total number and spatial variation in undetected roosts; (4) test whether our predictions of bat roost density improved explanations of the locations and intensity of VBR outbreaks in livestock; and (5) use the inferred projection of VBR outbreaks to re-assess the burden of rabies spillover.

## Materials and methods

### 1. Study area

The study area comprised three regions in southern Peru: Ayacucho, Apurimac, and Cusco (hereafter ‘AAC’, joint area = 136,697 km^2^, Fig. 1). AAC is largely composed of inter-Andean valleys, with elevations ranging from ∼280 to ∼6,000 m above sea level. To avoid modelling high-elevation areas that are known to exceed the physiological tolerance of vampire bats, we excluded areas above 4,000 m from the study area (Fig. 1B) [23]. The smoothr package in R was used to smooth the resulting landscape, which contained complex boundaries with small holes and sharp corners [29].

**Fig. 1:**
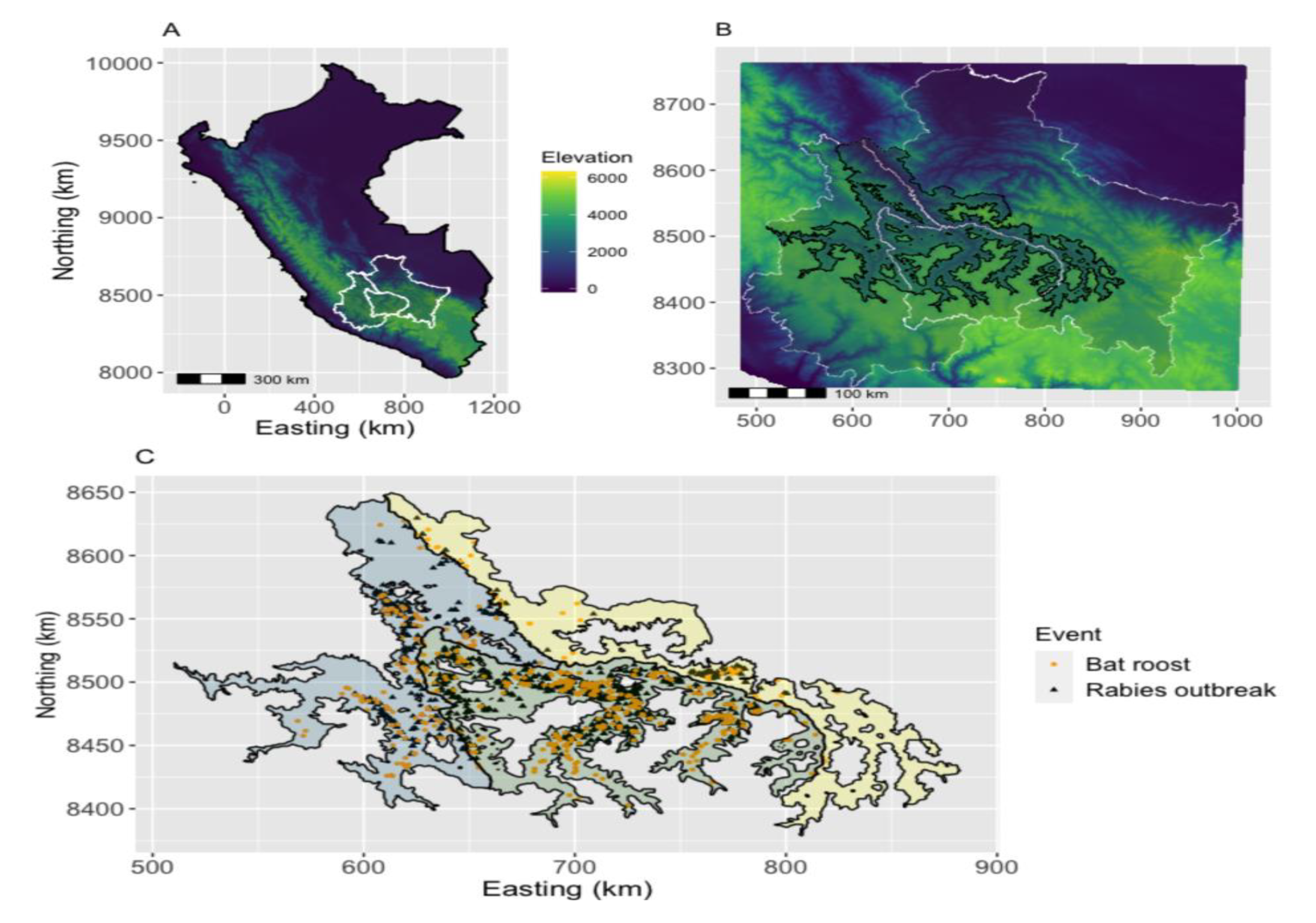
The study area with detailed spatial locations of vampire bat roosts and rabies outbreaks. A - Elevation in Peru (source https://www.worldclim.org) and the boundaries (white) of the departments (from left to right) of Ayacucho, Apurimac, and Cusco. B – Zoom to the three departments and the study area, the inter-Andean valleys (‘amebae shape’). These valleys exclude landscape with elevation above 4,000 m. C - The inter-Andean valleys of the AAC area (Ayacucho in light blue, Apurimac in light green and Cusco in light yellow), with the number of observed vampire bat roosts (n=563) and the number of reported rabies outbreaks between 2003 and 2021 (n=1,212).

### 2. Data on vampire bat roosts

We used the geolocations of 563 *D. rotundus* roosts in the AAC, collected between 2007 and 2021 (Fig. 1C). Roosts included natural (68%) and human-made structures (32%) but excluded foraging locations. Most natural structures inhabited by bats were caves (90%) with remaining classified as trees (5%) or ‘other’ (5%). Roosts in human-made structures were abandoned houses (47%), mines (24%), tunnels (15%), and ‘other’ (14%). Data originated from three sources: internally from the research group (2.4% of observations), from the Regional Government of Apurimac (22%, derived from a 2- year rabies control campaign [2014-2016], which included active searching for roosts), and from the National Service of Agrarian Health (SENASA, 75.6%). We analysed unique roost locations from these three partially overlapping datasets.

Apurimac had the highest roost detection effort because searches were carried out by both national and regional governments; moreover, our field studies in Apurimac initiated in 2007 (versus 2014 in Cusco and Ayacucho). Although national data from SENASA are more spatially extensive and represent information accumulated over the years, regional data were compiled using similar methods and may have been collected by the same individuals in some cases. Coordinates of roost locations were projected in Universal Transverse Mercator (UTM) and measured as northings and eastings in kilometers.

### 3. Environmental and anthropogenic variables

We selected environmental variables hypothesized to affect *D. rotundus* roost establishment, including (1) climate (annual mean temperature and annual mean precipitation); (2) topography (elevation, slope, and terrain rugosity index); (3) landcover (percentages of crop cover, tree cover, and permanent water); and (4) characteristics related to bat foraging (distance to the nearest river, prey density [a layer of cattle density and a layer of all livestock species density], number of rural population settlements per 2 km^2^ and human footprint). Table S1 (SI Appendix) provides the source and initial resolution of all variables. Data were collated as raster maps, set with the same extent (AAC area) and projection (UTM) (SI Appendix Fig. S1). Variables expected to act at a local scale such as climate or land cover, were rescaled to the finest resolution of all the variables (∼100 m). Variables expected to influence roost density at a broader scale (i.e., bat foraging) were used at their original resolution (1 or 2 km) or, in case of prey density, rescaled at 5 km resolution (the original resolution was 10 km). Prior to model fitting, variables were standardised (shifted to zero mean and scaled to unit variance) and the correlation between variables was assessed. To decrease the correlation between elevation (0-4,000 m) and other environmental variables, the layer of continuous elevation was reclassified as a binary variable defining suitable and less suitable altitudes for *D. rotundus* considering the threshold of 3,600 m indicated by previous research on the distribution of VBR in the AAC [23,30].

### 4. Modelling approach

To explain the heterogeneous and partially observed process of *D. rotundus* distribution, roost locations were treated as an inhomogeneous point pattern that follows a log Gaussian Cox process (LGCP) with intensity λ(*s*) at coordinates *s*. An LGCP is double-stochastic, because it is a hierarchical combination of a Poisson process at the first level (where the locations of the points are conditionally independent) and a Gaussian random field (GRF) at the second level (to account for spatial correlation in the data) [31,32]. We selected an LGCP to take advantage of point locations to explain the intensity process that creates the observed roost distribution.

Gaussian random fields are spatially continuous structured random processes that typically have dense covariance matrices that are computationally demanding, requiring long run times. An efficient way to introduce these structures in the model is to approximate the continuously indexed GRF by a spatially tessellated approximation to a Stochastic Partial Differential Equation (SPDE) [32,33]. Our model framework was developed using the Integrated nested Laplace approximation (INLA) joint with the SPDE approach, implemented via the inlabru R package [34]. Model fitting and inference were carried out in R version 4.1.2, using the packages R-INLA version 22.04.09 and inlabru development version 2.6.0.9003 [34–36]. We developed two main models that we present below: a detectability model to account for uneven observation effort in the study area, and the model of expected roost density, which was informed by the detectability model.

#### Detectability model

To ensure the patterns identified by our model were not observation artefacts and to robustly estimate the total number of roosts, it was necessary to correct for the uneven observation effort. Because data on the number of person-hours spent searching for roosts was unknown, we used the spatial accessibility of each location to approximate search effort. We approached this as a pre-analysis step, akin to some distance sampling analyses that fit a detection function separately from the ensuing habitat modelling [37]. Specifically, the detectability model described roost detections as a flexible function of accessibility to the main source of roost detection, measured as the mean travel time separating each grid cell from any of the 7 SENASA offices (SI Appendix, Table S1) [38]. To distinguish between regions that had different overall effort and, to include the effort from the Regional Government of Apurimac, we defined region as a two-level factor covariate separating Apurimac (highest effort) from Ayacucho and Cusco (combined). The observation model is defined by the expression:

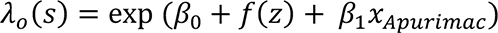

where *λ*_*o*_(*s*) is aimed to capture detection probability at location *s*, *β*_0_ is the intercept term, *f*(*z*) is the non-linear effect of the accessibility layer modelled using a random walk model of order 1 latent effect, and *β*_1_*x*_*Apurimac*_ represents the effect of region *x* (Apurimac is the reference). The predicted posterior mean derived from the observation model was normalised by assigning the probability 1 at the origin, which implies that a roost at the location of the SENASA offices is guaranteed to be known, resulting in the spatial probability of roost detection.

#### Matérn correlation model

The Matérn correlation model explains the scale of spatial dependency between neighbouring locations. This correlation structure is expressed over a discretisation of space known as the mesh (SI Appendix Fig. S2), a finite grid of triangulations of the spatial domain that approximates smooth random effects within the model [39]. Here, the mesh was defined within the smoothed boundaries of the inter-Andean valleys of the AAC area (the domain), and the resolution of the mesh was driven by the observed roost locations (hence directing more modelling detail in areas with detections). The correlation structure was defined according to the following equation:

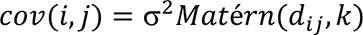

where the covariance between any two locations depends on their distance (*d*), on the range (*k*) of the Matérn function and on the spatial variance (σ^2^). This model included Penalised Complexity-priors (PC-priors) for range (*k*, practical range) and sigma (σ, the marginal standard deviation) [40].

#### Roost model

The expected density of vampire bat roosts (*λ*(*s*)) was fitted as a function of an intercept (*β*_0_), variables for roost distribution (*X*(*s*)), a spatial random field (*f*(*s*)), and a spatial offset (*ε*). The offset is the probability of roost detection from the detectability model, to adjust for the uneven observation effort. The roost model is defined according to the following expression:

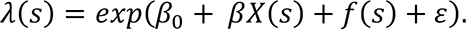

We used a forward addition procedure of variable selection, based on model conditional predictive ordinate (CPO), deviance information criteria (DIC), and Watanabe-Akaike information criteria (WAIC) to identify the most parsimonious model. If two variables were correlated (Pearson’s correlation, r > 0.6), the variable that better explained the response was retained, while the other was excluded from the model. To explore competition between bat colonies, as this behaviour is sometimes observed in other colonial species (e.g., seabirds), we considered adding the possibility of repulsion between roosts; however, the pairwise Euclidean distances between roosts demonstrated no evidence of such an effect (SI Appendix Fig. S3) [41].

#### Spatial predictions and estimates of abundance

To predict counts of roosts from the LGCP (i.e., an LGCP originates predictions in continuous space), the mesh was converted into 8-km^2^ grid cells, a dimension based on mesh’s resolution. We then used the fitted LGCP model to predict the number of roosts in each grid cell, generating samples from the posterior of all model parameters. To integrate the SPDE effect into roost count prediction, we projected integration weights to the mesh nodes and replaced the integral with a weighted sum in each grid cell. We predicted both the detected roosts (not correcting for the uneven effort) and the total expected roosts, in the latter case setting observation effort in all cells to the maximum such that observation probability was uniform. The posterior distribution of undetected roosts was calculated by subtracting the number of observed roosts (i.e., 563) from the density of the expected total count. We then estimated the posterior distribution for the total count that encompassed systematic stochasticity and uncertainty in parameter estimates, with and without adjusting for uneven effort.

#### Model validation

We validated predictions of roost expected density within our study area by refitting the model to a randomly selected 70% of the grid cells and reserving points falling inside the remaining 30% of the cells as a validation set (SI Appendix Fig. S4). We computed the distance between predicted and observed number of roosts in the validation set as a pseudo-R^2^ for count data [42]. Although discretising space into a regular grid required some subjectivity in choosing the size of the discretised units, the grid size was defined based on the mesh.

### 5. Explaining the distribution of past VBRV outbreaks

We hypothesized that our predicted roost distribution would improve understanding of the distribution of past livestock rabies outbreaks (i.e., defined as a single spillover from bats to livestock), identify high risk areas for future outbreaks, and identify locations with excess underreporting of outbreaks. For this purpose, we fitted a generalised linear model to the number of VBRV outbreaks that occurred between 2003 and 2021 (1,212 outbreaks, Fig. 1C and Fig. S5). We used a negative binomial likelihood to account for the overdispersion in the counts of rabies outbreaks in each 8-km^2^ grid cell. The model included an offset describing the probability of reporting rabies outbreaks in each grid cell due to accessibility to the SENASA offices [17]. We compared the performance of four alternative measures of bat distribution: (1) the observed number of roosts in each grid cell (raw data); (2) a kernel smooth of the raw data to represent a naive interpolation of bat density based on observed data without covariates or bias correction; (3) the predictions of our LGCP model corrected for the uneven effort; and (4) a smooth of these predictions (hereafter, Bat Utilization Distribution [BUD]), using the same parameters as the smooth for raw roost data. In addition to one of the four measures of bat distribution, we included the number of months since the first rabies outbreak occurred in each district to account for the geographic expansion of VBRV across AAC. We also included cattle density, reasoning that rabies would be more detectable in areas with more cattle [13,23,43]. We assessed the goodness of fit of eight competing models using Akaike information criterion (AIC) and model deviance pseudo-R^2^. We used the best model to predict the spatial distribution of past rabies outbreaks with and without correcting for accessibility to reporting centres. To assess the burden of rabies spillover, we estimated the underreporting factor as the ratio between predicted rabies outbreaks (corrected by uneven accessibility) and the reported rabies outbreaks in each grid cell. This analysis was restricted to grid cells that had at least one reported rabies outbreak to exclude predicted outbreaks in areas that as of 2021, remained rabies free (i.e., areas with high future risk).

## Results

### Detectability and landscape correlates of vampire bat roost establishment

The detectability model showed that roost observation decayed in a sigmoid shape with increasing travel time to SENASA offices. The inflection point of the sigmoid indicated high roost detectability up to 55 minutes of travel, with detection probability approaching zero for roosts that were more than 7 h from any SENASA office (Fig. 2A). The probability of detecting a roost increased with landscape accessibility and in areas that had more effort, particularly in Apurimac (Fig. 3A). Consistent with expectations for bat home range, the correlation structure of the Matérn model showed strong clustering of roosts up to distances of 10 km (half scale dependency) and positive correlations up to 30 km (Fig. 2B).

**Fig. 2:**
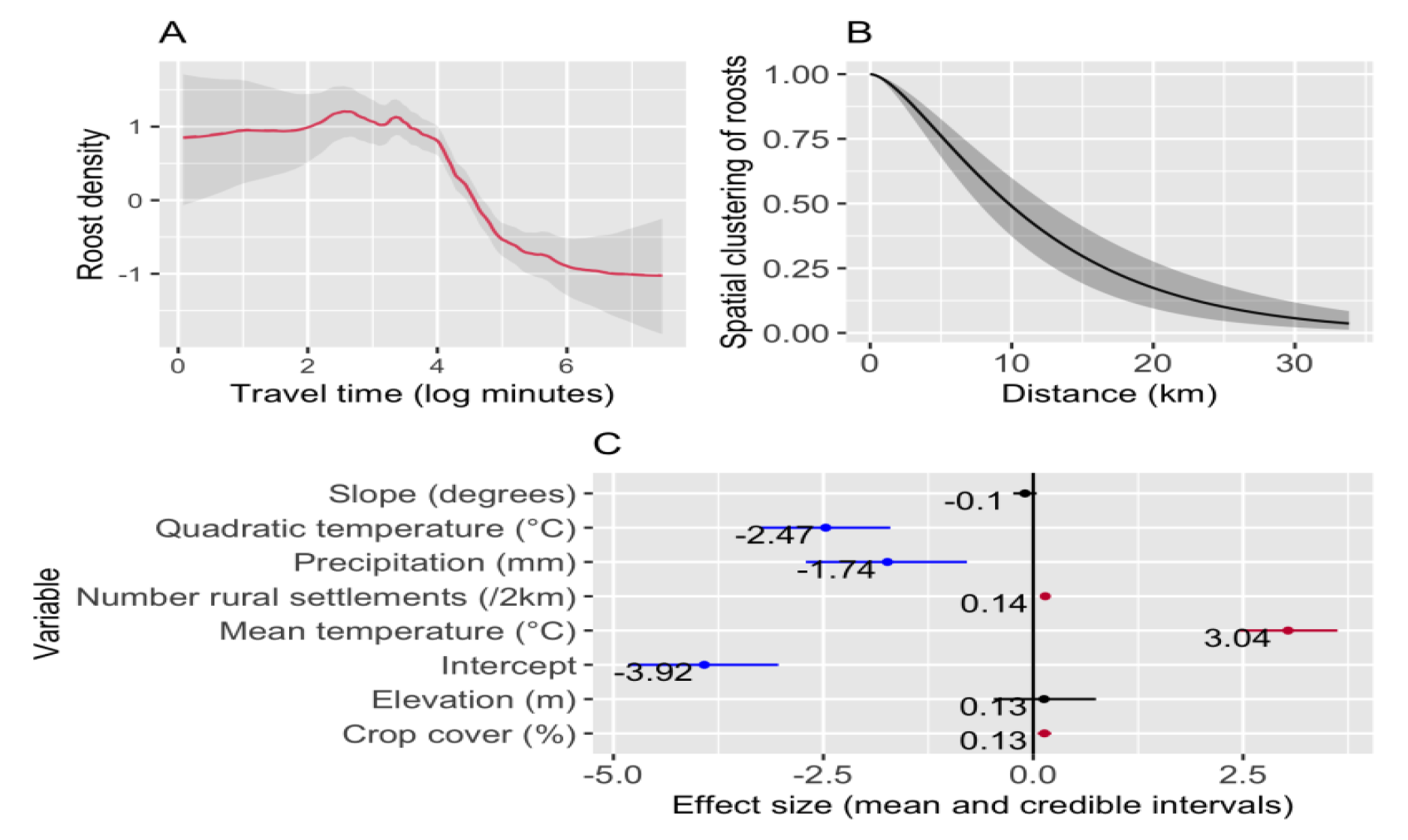
Factors affecting roost detectability and expected density: A – Non-linear effect of accessibility. X-axis is travelling time (log minutes, i.e., exp (4) = 55 min, and exp (6) = 7 h) and Y-axis is the mode of roost density (in the linear predictor scale). B - Range of the Matérn correlation model showing the mean correlation (Y-axis) in function of the distance between points. As the correlation range is on the log-intensity scale, on the intensity scale the range of correlation is shorter. Both in A and B, the grey shaded areas show the 95% credible interval. C - Posterior mean and respective 95% credible intervals of fixed effects of roost density. Red for positive significant coefficients (i.e., positive credible intervals), black for non-significant coefficients (i.e., include 0), and blue for negative significant coefficients (i.e., negative credible intervals).

**Fig. 3:**
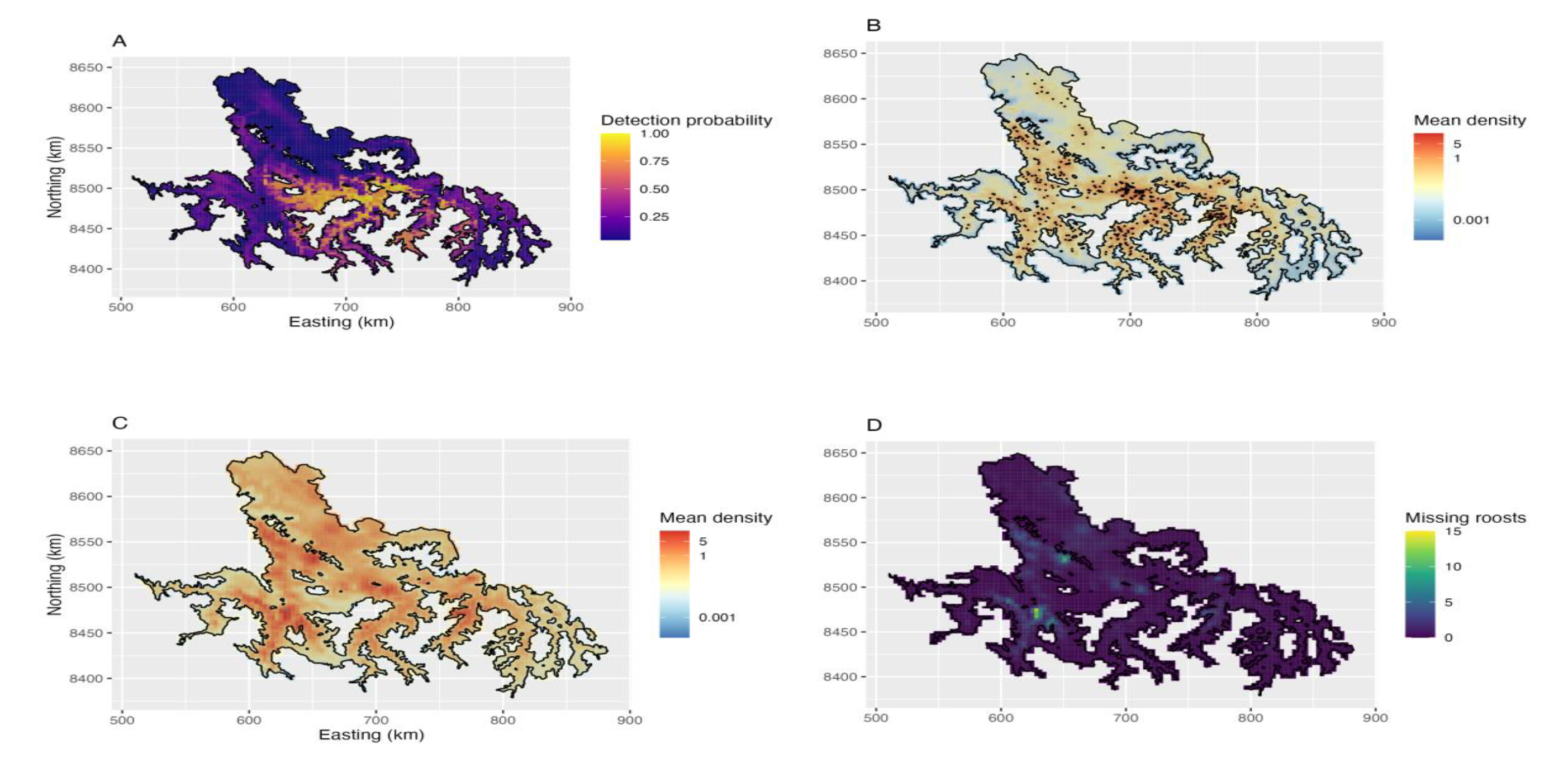
Spatial predictions of the detectability and roost models. A - Probability of detecting one roost (output from the detectability model). The detection probability is a combination of accessibility and surveillance effort in each region. Yellow for high probability of roost observation and dark purple for low probability of roost detection. B - Predictions of detected roosts. The black points are the 563 observed roosts. C – Predictions of the expected roosts when adjusting for the uneven effort. In both B and C, the predicted posterior mean of roosts was mapped; also, maps have the same scale, and the same continuous colour gradient was applied, with red indicating a higher mean of predicted roosts. D – Missing roosts (C-B). Light colour indicates higher number of missing roosts.

*D. rotundus* roost density was strongly influenced by climate, increasing with mean temperature until an optimum of 16.4 °C (Fig. S6) and decreasing in areas with higher average precipitation (Fig. 2C). Roost density also increased in proximity to rural human settlements and in areas with high crop cover, possibly both reflecting the availability of livestock prey. Terrain topography had marginal but non-significant effects on roost density with more roosts in areas with optimal elevation (<3,600m) and lower slopes (Fig. 2C).

### Distribution and density of *D. rotundus* roosts

The spatial projection of expected roost density resembled the observed roost distribution (pseudo-R^2^ = 0.60; Fig. 3B) and predicted a similar number of roosts as those used in model fitting (observed: 563 roosts; predicted: mean = 579, credible interval [CI]: 529-628; Fig. S7A). Pseudo-R^2^ decreased to 0.36 when removing the spatial random field, indicating that the GRF captured variability in roost density not explained by the environmental covariates. In our 30% hold out validation data, the full model with environmental covariates and the spatial random field explained 47% of the variance in roost density. Correcting for uneven observation effort revealed a total of 2,358 roosts (CI: 2,136- 2,684), implying that 1,795 roosts (76%) remain undiscovered (Fig. S7A). Importantly, roost detectability varied considerably over space, with as few as 4% of roosts known in some areas. Further, undiscovered roosts were spatially clustered, with hotspots identified in western and northern parts of our study area (Fig. 3C-D). The predictions of the posterior distribution for the total count of roosts had higher uncertainty when adjusting for effort, as it predicts the total abundance expected in the AAC area, accounting for observed and missing roosts (Fig. S7B-C).

### Explaining the distribution of past VBRV outbreaks

In univariate models of the alternative descriptions of the vampire bat distribution, the smoothed predicted roost distribution (BUD) was the strongest predictor of rabies outbreaks, explaining 22% of the distribution of past rabies outbreaks (Table 1). When adding the additional covariates describing the geographic expansion of VBRV across AAC and cattle density to the BUD model, explanatory power increased by a further 15% (Full model pseudo-R^2^ = 0.37). Alternate models using other versions of the bat distribution and the additional covariates had similar pseudo-R^2^ (0.31–0.36), but consistently higher AIC (ΔAIC=128.4–29.6, Table 1). Spatial predictions of rabies outbreaks from the best model (model 1), generated with and without correcting for accessibility to reporting offices (Fig. 4A-B), revealed strong spatial heterogeneity, signaling probable hotspots of unreported rabies spillover across our study area or high-risk areas where future outbreaks might go unreported. Comparison of predicted to observed rabies outbreaks showed that, on average, the number of rabies cases in livestock may have been 7.1 times (95% CI: 3.8–10.5) higher than officially reported (Fig. 4C).

**Fig. 4:**
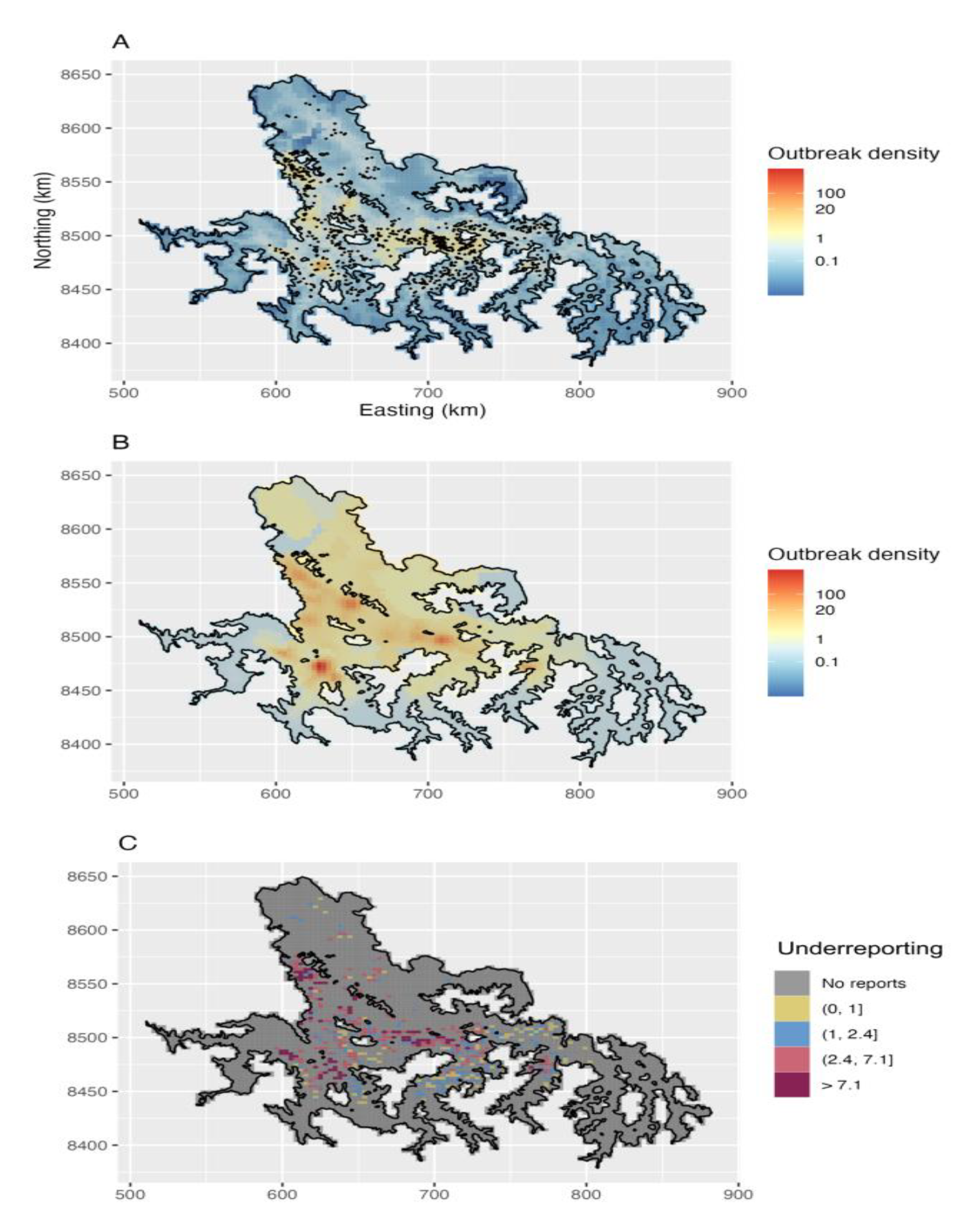
Spatial predictions of rabies outbreaks and heterogeneity in underreporting. A – Predictions of reported rabies outbreaks. The black points are the 1,212 reported rabies outbreaks in livestock between 2003 and 2021. B – Predictions of rabies outbreaks after correcting for accessibility to reporting offices. A and B have the same scale. A continuous colour gradient was applied, with red indicating a higher number of outbreaks. C – Underreporting in cells where outbreaks were reported. The cells were grouped according to the ratio between predictions corrected for accessibility and reported outbreaks. The two thresholds used were the average (i.e., 7.1) and the median (i.e., 2.4) of underreporting in the cells where outbreaks were reported (493 cells out of 4,797).

**Table 1:**
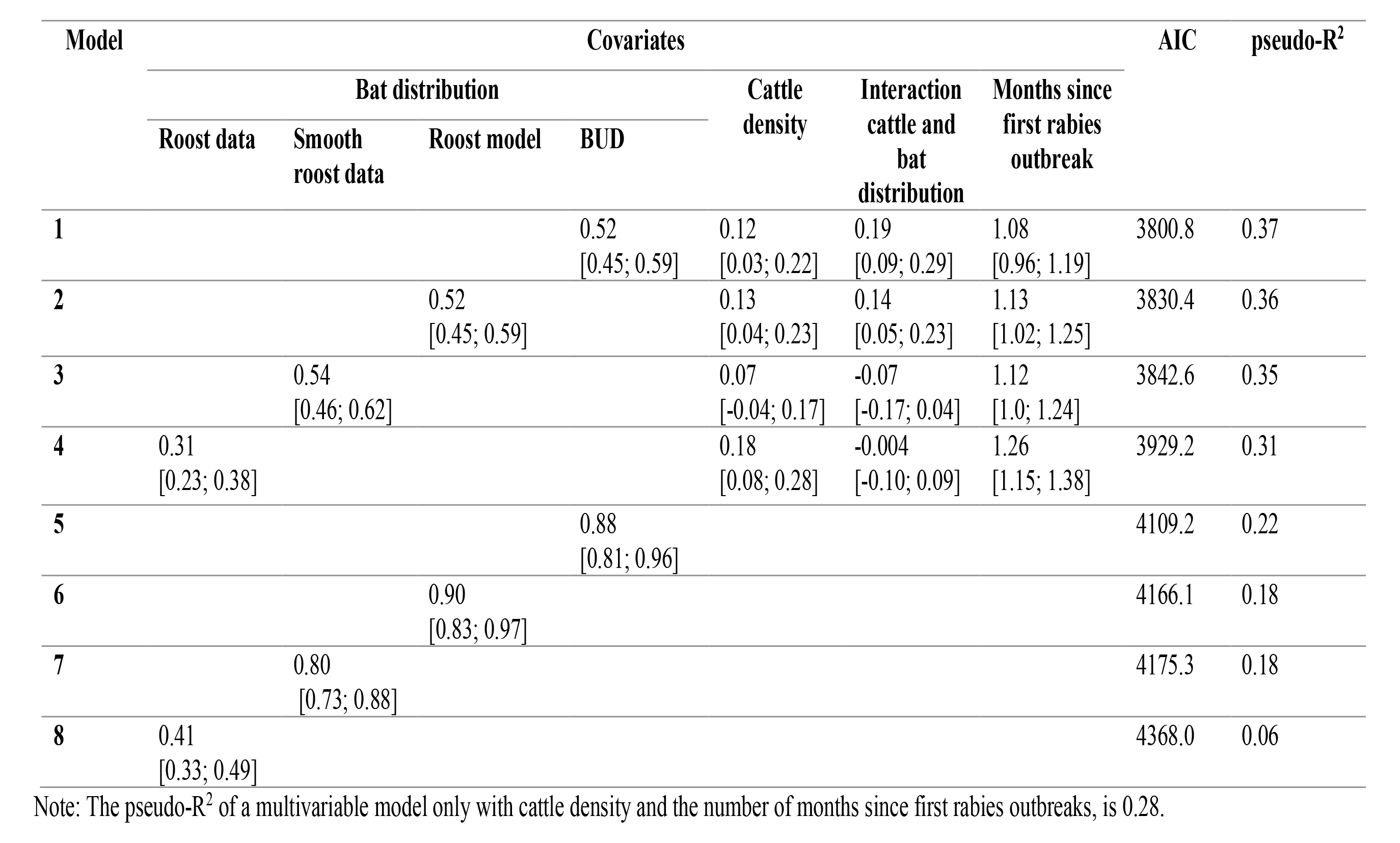
Estimate and 95% confidence intervals of the covariates included in the eight models used to compare the performance of four measures of bat distribution. The estimates of the fixed effects are presented in the logarithm and standardised forms. Models are displayed in descending order, from higher to lower AIC and pseudo-R^2^.

## Discussion

Understanding the spatial distributions of wildlife reservoirs of zoonoses is vital to anticipate and prevent spillover but undermined by incomplete, biased, and spatially autocorrelated observations of animal locations. Focusing on an important bat reservoir in Latin America, our study provides a transferrable statistical approach to infer the spatial distribution of partially observed wildlife [7]. In our study, spatial heterogeneity in vampire bat colonies was associated with topographic, ecological, and anthropogenic factors but also strongly affected by previously unrecognised observation biases. Indeed, our results indicate that only 24% of vampire bat roosts are known to authorities and models correcting these biases revealed unrecognised hotspots in vampire bat distribution. Finally, predictions of bat density improved models of the locations and intensity of historical rabies outbreaks, revealed spatial heterogeneity in the reporting of rabies outbreaks, and increased the projected burden of rabies mortality. By identifying spatial hotspots of bat roosts and potential future rabies outbreaks, our results may improve the allocation of limited resources for vampire bat management and rabies prevention.

Our model indicated that temperature and precipitation limit the formation of vampire bat roosts, corroborating results from earlier analyses conducted across the geographic range of this species [26–28]. Although roosts of *D. rotundus* require more than 45% humidity, the mechanism by which precipitation acts on vampire bat distribution is poorly understood [44]. The negative effect of precipitation in our model is most likely explained by the increase in observed bat roosts from the rainy north to the dry climate in the centre and south of the valleys [45]. Temperature effects were expected as *D. rotundus* is a poor homeotherm and environmental temperatures below 10°C or above 37°C are not tolerable [46,47]. Of note, however, the optimal environmental mean temperature found in our study (i.e., 16.4°C) is relatively low compared with the recommended temperature for maintaining vampire bats in captivity, between 21°C and 27°C [48]. It is possible that this difference reflects our use of ambient temperature in our model, rather than within roost temperature measurements, which were not measured in our study. Vampire bat roosts would generally be expected to have more stable and lower temperatures than ambient, particularly in daytime hours when bats occupy roosts, but this relationship may vary depending on local conditions and roosts characteristics. Further, AAC valleys represent a range limit for this species in southern Peru with higher elevation areas farther south experiencing intolerable conditions. Assuming the recommended captive maintenance temperatures are valid, our results indicate that in the AAC area, vampire bats exist in sub-optimal temperatures. This implies the possibility of physiological trade-offs that influence the susceptibility to or tolerance of vampire bats to infection. For example, female tree swallows (*Tachycineta bicolor*) persisting in suboptimal sites in Alaska had evidence of impaired immunity and populations of whistling tree frogs (*Litoria verreauxii verreauxii*) in low quality habitats were less likely to persist through epizootics of chytridiomycosis [49,50]. For vampire bats, heightened susceptibility to lethal rabies infection is predicted to have large dynamical consequences on viral prevalence and spillover [21]. If impaired immunity in these populations is verified, we speculate this might contribute to the disproportionate burden of rabies in the AAC, which despite its small size, accounts for most rabies outbreaks at the national level [23]. It is also conceivable that the high genetic divergence of AAC vampire bats from those in tropical areas included physiological adaptations to low temperatures which may have unpredictable trade-offs with immunity [30,51,52].

We also found effects that are consistent with the influence of human activities on vampire bat roost establishment. For example, the negative effect of slope may reflect use of artificial (human-made) roosts, which are generally constructed in flatter areas. The positive effects of the number of rural human settlements and crop cover on vampire bat roost establishment likely reflect the availability of livestock, which are the main food source for *D. rotundus* [43,53,54]. Although it is unexpected that livestock itself was not retained by the model, we suggest that the global layers used, while suitable to access broad scale effects of livestock, are less representative of variation in livestock density at fine spatial scales than the more finely resolved measures of human presence [43]. As such, our findings support the hypothesis that anthropogenic activities favour vampire bat roost establishment and may facilitate spatial expansions of this species to new areas within climatically tolerable regions [30].

Despite the influence of environmental covariates in explaining vampire bat roost density, the higher pseudo-R^2^ after adding the GRF (pseudo-R^2^ = 0.36 versus 0.60) indicated that the spatial dependency among roosts explained considerable variation in roost density. For example, in cranes and seabirds, such dependencies may arise from dispersal limitations, inter-specific competition, disturbance, or social interactions [10,11,31]. Here, social interactions may be particularly important to explain spatial dependency at the roost level, as vampire bats are social species and often use multiple roosts, mostly within small areas (2 to 3 km radius) [55,56]. In the AAC, correlation between roosts remained positive up to 30 km (Fig. 2B). The spatial extent of this clustering is somewhat unexpected given vampire bat home ranges are believed to extend only 5–10 km. It is conceivable that longer distance connectivity – occasionally reported – may be more common than previously recognised, which would have important implications for rabies spread [44,47,53,57]. Our results also corroborate the absence of competition between vampire bat roosts, which would have been expected to generate a negative correlation at small distances [31]. More generally, our results illustrate the value of accessing the shape of the Matérn correlation range to understand and generate hypotheses about animal ecology and social behaviour.

Besides spatial dependency, observation biases are expected in most natural habitats but have not historically been incorporated into distribution modelling for wildlife. Our observation model captured a decay in vampire bat roost detection as accessibility to reporting offices decreased (Fig. 2A) and a sharp decrease in observed roosts in Ayacucho and Cusco (Fig. 3A). When correcting predictions for the uneven effort, our model estimated that the number of undetected roosts was more than triple the number of known roosts. Moreover, missed roosts were spatially clustered, rather than uniformly spread across the study area. Of note, some hotspots of missing roosts are currently in rabies free areas that have neighbour areas with current viral circulation, indicating high risk of rabies spread into areas where outbreaks in bats and thus livestock might rapidly ensue [23]. This highlights an important outcome of incorporating observation processes into model projections, enabling interventions such as preventive bat culls or livestock vaccination that might prevent spillover or reduce its burden in high-risk areas. More generally, improved understanding of the true spatial distribution of vampire bats facilitates spatially synchronized control, which is predicted to be central to successfully managing rabies in endemic areas, but until now, it was precluded by gaps in our knowledge of bat distribution [21,43,58].

In addition to identifying areas at high risk of rabies outbreaks, our predictions of detected and undetected roosts improved understanding of the spatial distribution of VBRV outbreaks in livestock and revealed hotspots of disease underreporting. Previous work carried out in Brazil, which assessed the implications of the geographic distribution of vampire bat roosts in the occurrence of rabies in livestock, demonstrated a positive correlation between detected bat roosts and density of rabies outbreaks in livestock [53,58]. Here, compared with raw roost data and its smooth surface, our roost distribution model had a better fit and explained more of the spatial pattern of past rabies outbreaks in Peru (Table 1). However, differences between the performance (i.e., pseudo-R^2^) of models with the smoothed raw data and roost predictions were relatively small. This may reflect the fact that roost detection and notification of rabies outbreaks were generated by similar observation processes, both being affected by distance from the nearest reporting centre. Nevertheless, the modelling approach presented here is novel and different from previous studies as we accounted for undetected bat roosts. This is particularly advantageous for identifying high-risk areas for disease spillover which can be targeted to strengthen rabies awareness (i.e., increasing chances of reporting), surveillance, and control (i.e., vaccination and synchronised culling, bat vaccination) [59].

In Latin America, rabies is a neglected zoonotic disease, and suspected outbreaks in livestock are notified via passive surveillance. Hence, underreporting of rabies outbreaks leads to underestimation of the true burden of the disease [12,17,54]. In previous research, Benavides et al. [17] used questionnaires to estimate underreporting in AAC, predicting that mortality from VBRV was 4.6 times (95% CI: 4.4– 8.2) higher than officially reported, and inferring spatial heterogeneity in underreporting at district level. Here, without use of extensive questionnaires, our model estimated that between 2003 and 2021, mortality from VBRV in AAC was 7.1 times (95% CI: 3.8–10.5) higher than officially reported. In addition, our predictions of rabies underreporting demonstrated spatial heterogeneity at a finer resolution than previously possible (Fig. 4C). These results have two distinct implications. First, our elevated estimate of underreporting implies an underestimation of the disease’s true burden which can be used to inform VBR management. Secondly, our ability to estimate underreporting at fine spatial scales empowers more precise geographic allocation of educational campaigns to encourage reporting where they are most needed.

Although the statistical approach presented here provides advances in modelling partially observed wildlife disease reservoirs, our projections may be improved in several ways. Future work could take advantage of more sophisticated extensions of SDMs that integrate distinct but complementary data on vampire bat occurrence (e.g., roosts and reports of bat bites) [31]. Additionally, integrating the observation and the roost models, rather than modelling them separately as done here, would also be advantageous to fully propagate uncertainty across models. A joint likelihood approach with a parametric form for the detection function would be a potential approach to extend our framework [60,61].

In conclusion, we present a transferrable statistical approach to model the spatial distribution of difficult to observe species, demonstrating the benefits of consider spatial autocorrelation and observation effort. To our knowledge, this is the first-time observation effort and spatial autocorrelation have been used to reconstruct the map of likely roosts for any bat species. Our outputs identified putative hot and cold spots of vampire bat roosts and risk areas for VBRV spread, and areas with high number of unreported rabies outbreaks. These results can be valuable in spatial models that explore determinants of viral maintenance and spillover risk and can guide effective monitoring of *D. rotundus* and prevention and control strategies for VBR, such as campaigns to increase awareness of rabies, preventive vaccination of livestock and in future, vaccination of vampire bat colonies. More generally, our results show how incorporating existing and routinely collected data into powerful statistical models can improve the management of zoonoses.

## Author contributions

Conceptualisation: R.R., J.M. and D.G.S.; Formal analysis: R.R.; Funding acquisition: T.E.R. and D.G.S.; Methodology: R.R., J.M., F.L., and D.G.S.; Resources: C.T., C.M.Z., W.V. and D.G.S.; Supervision: J.M. and D.G.S.; Visualization: R.R.; Writing-original draft: R.R., J.M. and D.G.S. R.R.; Writing-review and editing: All authors reviewed the and gave final approval for publication.

## Acknowledgments

The authors would like to thank Andrew Seaton (University of Glasgow) for assistance with the process of predicting counts from an LGCP; Luca Nelli (University of Glasgow) for the suggestion of relevant variables; all colleagues and students who provided valuable comments on the model, results, and draft: Iosu Paradinas (Asociación Ipar Perspective), all members of the <u>Streicker lab</u> (Nardus Mollentze, Laura Bergner, Jocelyn Pérez Lazo, Max Farrell, Megan Griffiths, Hollie French, Haris Malik, Alice Broos, and Matthew Arnold), Jorge Osorio (University of Wisconsin-Madison), Zulma E. Rojas-Sereno (Universidad Andres Bello), from the University of Glasgow: Yacob Haddou, Jana Jeglinski, and Landry Green, and Fergus Chadwick (BioSS) and Crinan Jarret (vogelwarte). Paulo Colchao and Manuel Sime for their support in roost detection. The dataset of VBRV outbreaks in livestock between 2003 and 2021 was kindly provided by SENASA.

## Data availability statement

The data and R scripts that support the findings of this study are available in Zenodo [62].

