## Supplementary material for "Incorporating environmental heterogeneity and observation effort to predict host distribution and viral spillover from a bat reservoir": SI Appendix

**Authors:** Rita Ribeiro[1], Jason Matthiopoulos[1*], Finn Lindgren[2], Carlos Tello[3,4], Carlos M. Zariquiey[3], William Valderrama[3,5], Tonie E. Rocke[6], Daniel G. Streicker[1,7*]

∗These authors share senior authorship

**Affiliations:**

[1]School of Biodiversity, One Health and Veterinary Medicine, College of Medical, Veterinary and Life Sciences, University of Glasgow, Glasgow, UK

[2]School of Mathematics, University of Edinburgh, Edinburgh, UK

[3]ILLARIY (Asociación para el Desarrollo y Conservación de los Recursos Naturales), Lima, Perú

[4]Yunkawasi, Lima, Perú

[5]Facultad de Medicina Veterinaria y Zootecnia, Universidad Peruana Cayetano Heredia, Lima, Perú

[6]U.S. Geological Survey, National Wildlife Health Center, Madison, Wisconsin, USA

[7]Medical Research Council - University of Glasgow Centre for Virus Research, Glasgow, UK

**Contents:**

S1 Spatial distribution and summary information of variables assessed in this study.

S2 Constrained refined Delaunay triangulation.

S3 Pre-analysis of the pairwise distance between vampire bat roosts.

S4 Model validation – data kept for model fitting and model validation.

S5 Number of rabies outbreaks in livestock in the inter-Andean valleys of Apurimac, Ayacucho, and Cusco between 2003 and 2021.

S6 Quadratic function of temperature.

S7 Posterior distribution of roost abundance.

**Disclaimer:** Any use of trade, firm, or product names is for descriptive purposes only and does not imply endorsement by the U.S. Government.

**S1 Spatial distribution and summary information of variables assessed in this study.**

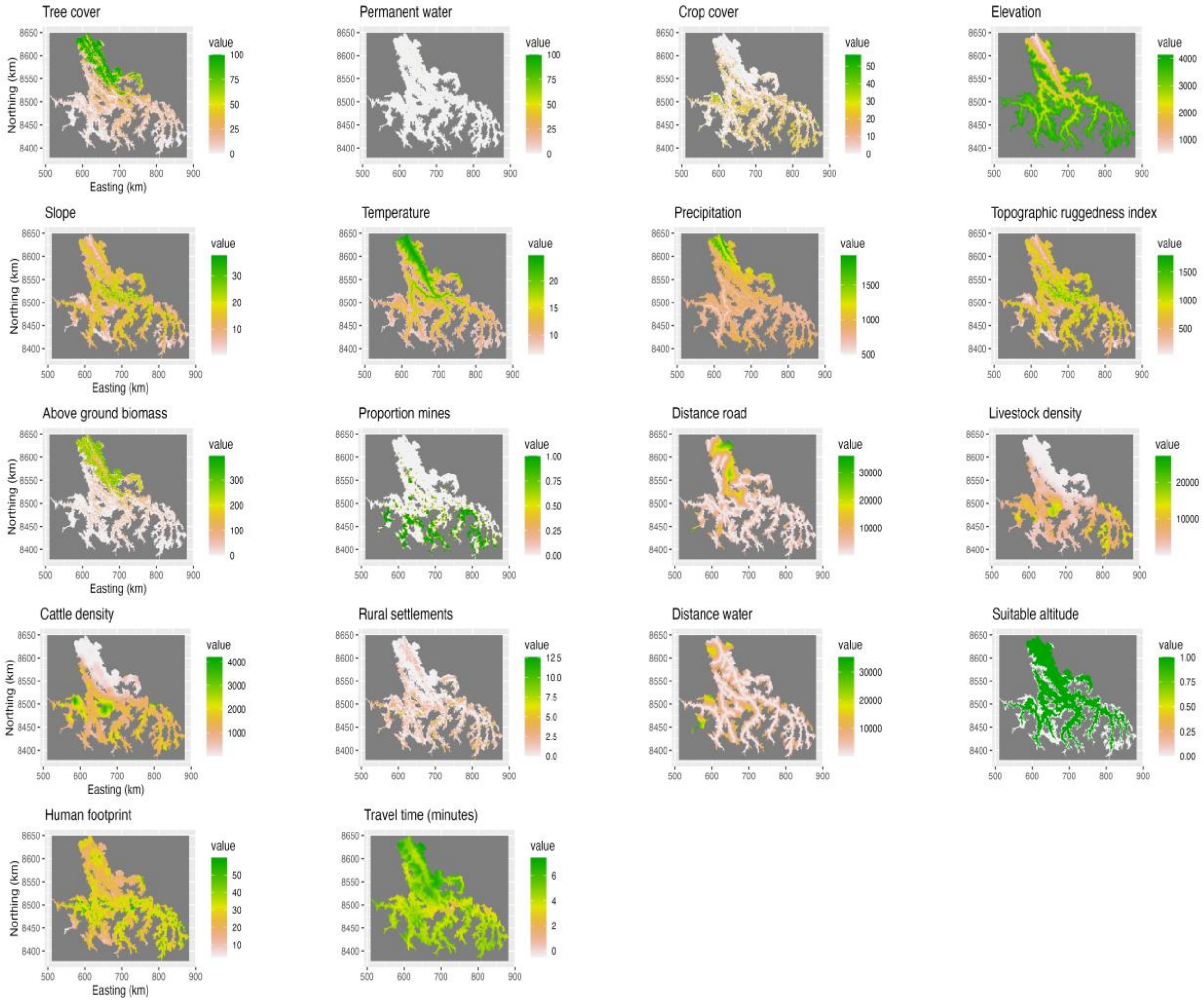


Fig. S1: Spatial distribution of the environmental variables assessed for the study in the inter-Andean valleys of Apurimac, Ayacucho, and Cusco.

Table S1: Environmental variables assessed for the study, original resolution, and source.

| **Variables** | | **Resolution** | **Source** |
|---|---|---|---|
| Climatic | Annual mean temperature (°C) | 30 seconds (or 0.0083 degrees ~ 1 km) | [1]<br>https://www.worldclim.org<br>Use getData() R package raster |
| | Annual precipitation (mm) | 30 seconds (or 0.0083 degrees ~ 1 km) | [1]<br>https://www.worldclim.org<br>Use getData() R package raster |
| Topographic | Elevation | 30 seconds (or 0.0083 degrees ~ 1 km) | [1]<br>https://www.worldclim.org<br>Use getData() R package raster; elevation derived from the SRTM elevation data |
| | Elevation reclassified to suitable and unsuitable for bat movement | 0.0083 degrees ~ 1 km | Derived from elevation. Binary covariate defining preferred and non-preferred habitats for *D. rotundus* considering the threshold of 3,600 m [2]. |
| | Slope | 0.0083 degrees ~ 1 km | Derived from elevation<br>Use terrain() R package raster |
| | Terrain ruggedness index (TRI) | 0.0083 degrees ~ 1 km | Derived from elevation using SpatialEco R library |
| Landcover | Percent vegetation cover for cropland land cover class | ~100 m | Copernicus, Proba-V [3] |
| | Percent vegetation cover for forest land cover class | ~100 m | Copernicus, Proba-V [3] |
| | Percent ground cover for permanent water land cover class | ~100 m | Copernicus, Proba-V [3] |
| | Above ground biomass (AGB) | ~100 m | The mass, expressed as oven-dry weight of the woody parts (stem, bark, branches, and twigs) of all living trees excluding stump and roots. [4] |
| | Mines | 1 km | A polygon layer of the mines present in the AAC area was extracted from https://geocatmin.ingemmet.gob.pe/geocatmin/ (Geological and mining cadastral information system from Peru), and used to create a raster with the proportion of mines per 1 $km^2$. |
| Host related | Cattle density | 5 arcminutes, ~ 10 km at the equator. | Gridded Livestock of the World (GLW3). Use of the dasymetric weighting version (DA) [5]. |
| | Animal density | 5 arcminutes, ~ | Gridded Livestock of the World (GLW3); resulted from the sum of cattle, horses, goats, sheep, and pig |

| | | | |
|---|---|---|---|
| | | 10 km at the equator. | density. Use of the dasymetric weighting version (DA) [5]. |
| | Number of rural population centres per 1, 2 and 5 km square | 1, 2 and 5 km | Data on the geographic locations of the rural population centres in the AAC area, and of the census population on those rural centres were obtained from two departments of Peru Government (INEI and CEPLAN) (https://www.inei.gob.pe) and https://www.ceplan.gob.pe/informacion-sobre-zonas-y-departamentos-del-peru/). This data was used to calculate the number of rural population centres per 1 $km^2$, 2 $km^2$ and 5 $km^2$. |
| | Human footprint | 0.0083 degrees (~1km) | https://sedac.ciesin.columbia.edu/data/set/wildareas-v2-human-footprint-geographic; [6]. |
| | Distance to the nearest river | 1 km | A polygon layer with the river network in the AAC area from https://mapcruzin.com was used to create a raster of 1 $km^2$ size, with the mean Euclidean distance to the nearest river. |
| Accessibility to SENASA offices (minutes of travel time) | | 0.0083 degrees (~1km) | Created from a global friction surface developed by Weiss et al. [7], and enumerating land-based travel speed. gdistance R package was used to create a transition layer and, with the points of the National Service of Agrarian Health (SENASA) offices, a layer of travel times.<br>The accessibility to the SENASA offices is represented by the travel time in minutes (logarithm scale) from each grid cell to the SENASA offices. |
| Region maps | Obtained from the GADM ( http://www.gadm.org/) database using the getdata function from the R raster package. | | |

**S2 Constrained refined Delaunay triangulation.**

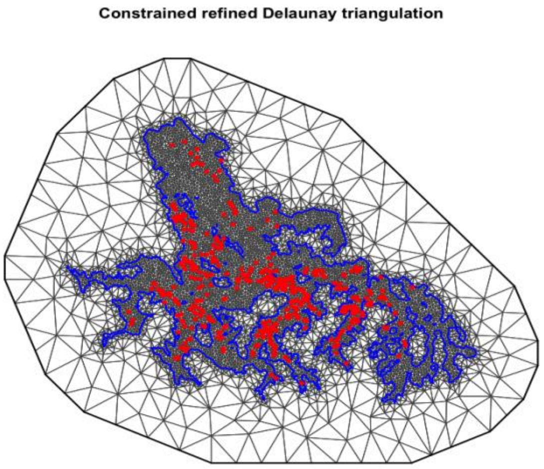


Fig. S2: Constrained refined Delaunay triangulation. The ‘mesh’ consists of 7,041 vertices, with small triangles with almost the same dimensions in the inner domain, where the predictions are important, and bigger triangles in the outer extension, to decrease the boundary effect. The blue line represents the smooth boundaries of the inter-Andean valleys of Apurimac, Ayacucho, and Cusco, and the red points are the 563 vampire bat roosts. The lengths of the mesh have the same units as the reference coordinate system.

**S3 Pre-analysis of the pairwise distance between vampire bat roosts.**

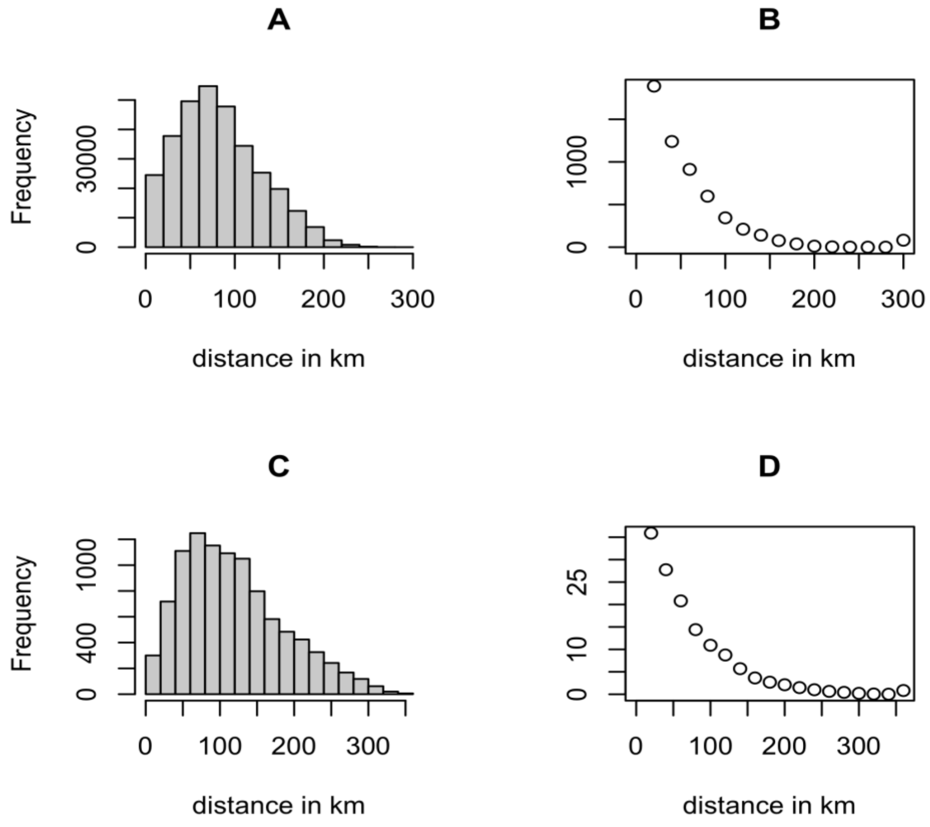


Fig. S3: Analysis of pairwise distance between roosts excludes hypothesis of repulsion between roosts. A - Histogram of the pairwise distance between the 563 roosts (units in km). B – Y-axis is the count of pairwise distances divided by each distance break and X-axis is each distance break of the histogram. C and D represent the same as A and B but for 100 simulated roosts within the boundaries of the inter-Andean valleys of the AAC area, to verify the relationship kept the same.

### S4 Model validation – data kept for model fitting and model validation.

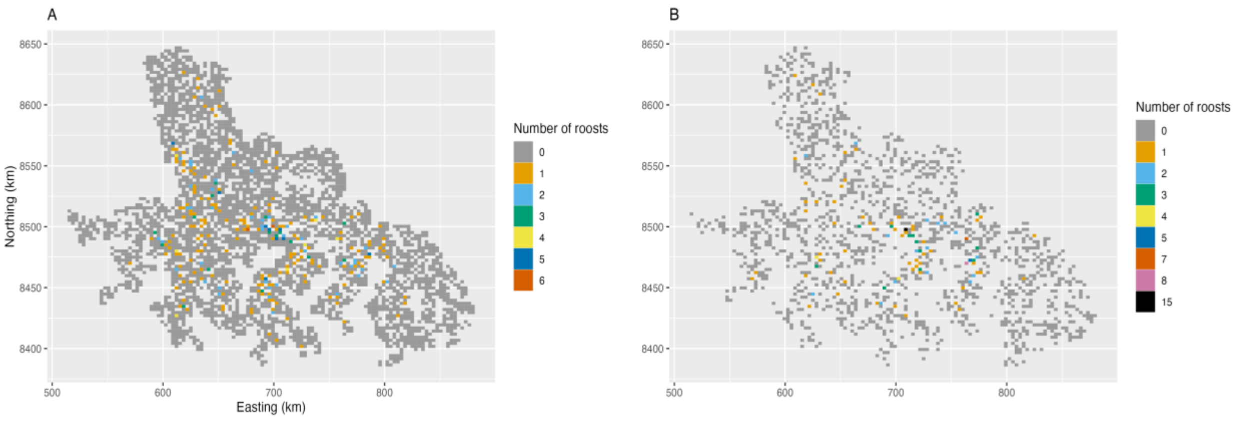


Fig. S4: Training and validation datasets for the roost model. A – 70% of the grid cells were randomly selected and kept as samplers for the model. Points falling on those cells were kept for model fitting. B – 30% of the cells removed after random sampling and kept for model validation. A discrete colour scheme was preferred to help with visualisation.

### S5 Number of rabies outbreaks in livestock in the inter-Andean valleys of Apurimac, Ayacucho, and Cusco between 2003 and 2021.

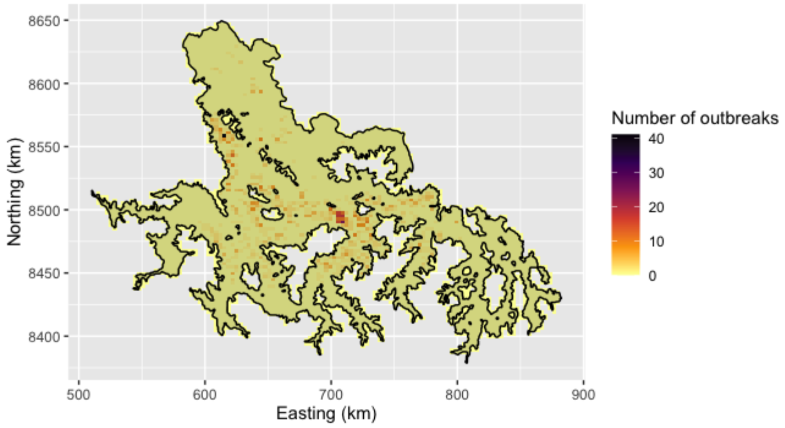


Fig. S5: Number of rabies outbreaks per grid cell between 2003 and 2021 (1,212 outbreaks). Darker colour means high number of outbreaks.

**S6 Quadratic function of temperature.**

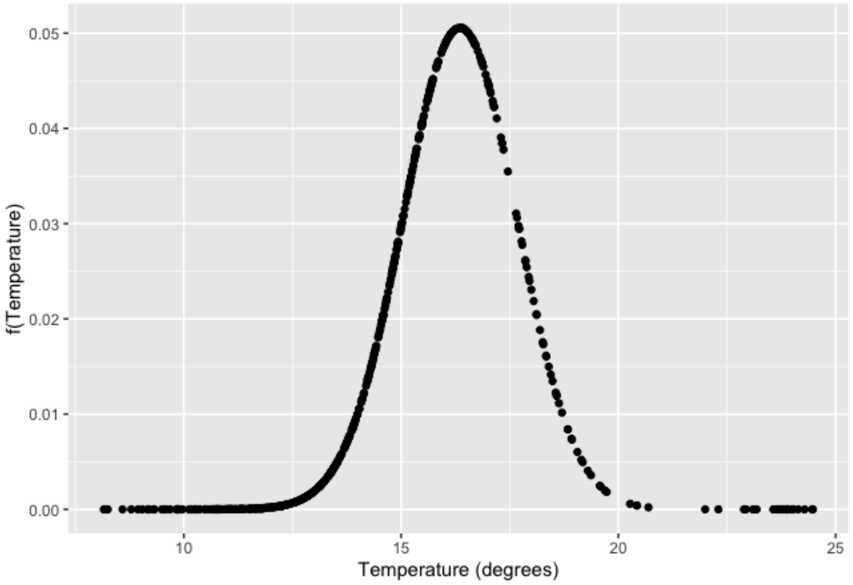


Fig. S6: Quadratic function of temperature. The quadratic function of temperature is derived from the expression f(*temperature*) = exp(*a***temperature_std*$^2$ + *b***temperature_std* + *c*). *Temperature_std* is the standardised temperature, *a* is the standardised coefficient for the quadratic term, *b* the standardised coefficient for the linear term, and *c* is the coefficient for the intercept. The optimal temperature (non-standardised temperature values) was computed based on: $X = ((-b)/2a) * \sigma + \mu$, where $X$ is the parabola vertex, $\sigma$ is the standard deviation of temperature in the non-standardized scale, and $\mu$ is the mean of temperature in the non-standardized scale.

**S7 Posterior distribution of roost abundance.**

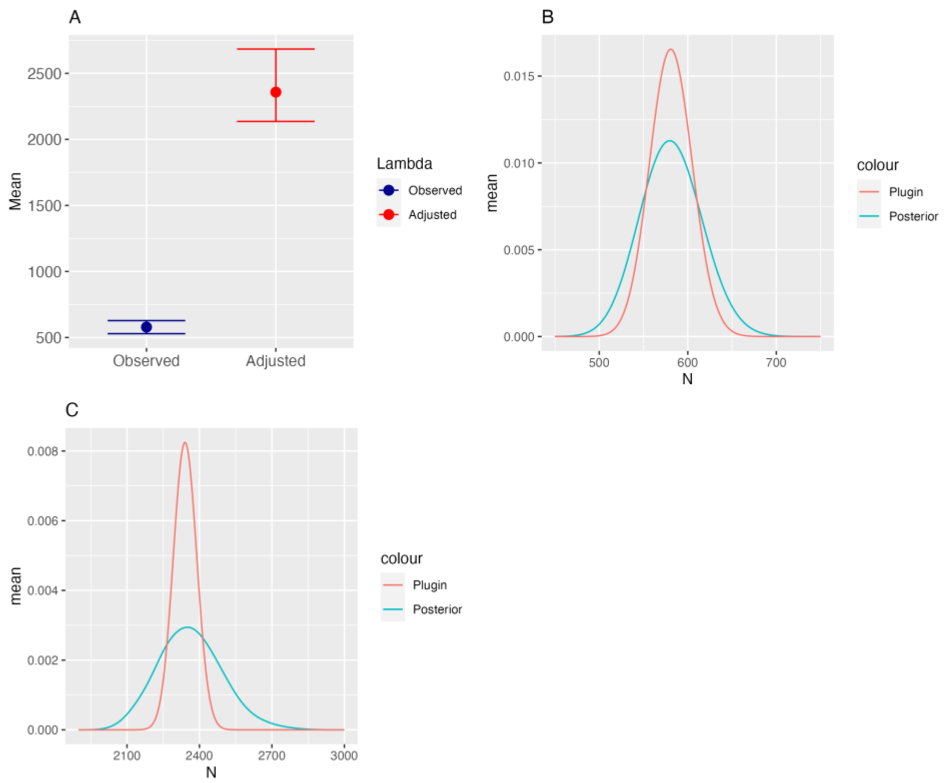


Fig. S7: Posterior distribution of roost abundance, corrected or not for observation effort. A – Predictions of the posterior mean of observed roosts ('Observed', blue dot), and predictions of total expected roosts ('Adjusted', red dot) and respective credible intervals (blue and red bars). B – Posterior distribution for the total observed roosts (not correcting for the uneven effort). C - Posterior distribution for the total expected roosts (correcting for the uneven effort. In both B and C, the 'posterior' encompasses systematic stochasticity and uncertainty in parameter estimate, whereas the 'plugin' only considers model stochasticity. We can interpret the difference between the two curves by the parameter uncertainty.
